# Global cataloguing of variations in untranslated regions of viral genome and prediction of key host RNA binding protein-microRNA interactions modulating genome stability in SARS-CoV2

**DOI:** 10.1101/2020.06.09.134585

**Authors:** Moumita Mukherjee, Srikanta Goswami

**Affiliations:** National Institute of Biomedical Genomics, Kalyani, West Bengal

**Keywords:** SARS-CoV2, UTR variants, RNA binding protein, microRNA

## Abstract

**Background:** The world is going through the critical phase of COVID-19 pandemic, caused by human coronavirus, SARS-CoV2. Worldwide concerted effort to identify viral genomic changes across different sub-types has identified several strong changes in the coding region. However, there have not been many studies focusing on the variations in the 5’ and 3’ untranslated regions and their consequences. Considering the possible importance of these regions in host mediated regulation of viral RNA genome, we wanted to explore the phenomenon.

**Methods:** To have an idea of the global changes in 5’ and 3’-UTR sequences, we downloaded 8595 complete and high-coverage SARS-CoV2 genome sequence information from human host in FASTA format from Global Initiative on Sharing All Influenza Data (GISAID) from 15 different geographical regions. Next, we aligned them using Clustal Omega software and investigated the UTR variants. We also looked at the putative host RNA binding protein (RBP) and microRNA binding sites in these regions by ‘RBPmap’ and ‘RNA22 v2’ respectively. Expression status of selected RBPs and microRNAs were checked in lungs tissue.

**Results:** We identified 28 unique variants in SARS-CoV2 UTR region based on a minimum variant percentage cut-off of 0.5. Along with 241C>T change the important 5’-UTR change identified was 187A>G, while 29734G>C, 29742G>A/T and 29774C>T were the most familiar variants of 3’UTR among most of the continents. Furthermore, we found that despite of the variations in the UTR regions, binding of host RBP to them remains mostly unaltered, which further influenced the functioning of specific miRNAs.

**Conclusion:** Our results, shows for the first time in SARS-Cov2 infection, a possible cross-talk between host RBPs-miRNAs and viral UTR variants, which ultimately could explain the mechanism of escaping host RNA decay machinery by the virus. The knowledge might be helpful in developing anti-viral compounds in future.

## Introduction

Outbreak of novel coronavirus SARS-CoV2 has been proclaimed as a pandemic by the World Health organization (WHO). From the first report of infection in Wuhan, China on December 31, 2019, the virus has infected 7.21M people worldwide till 9th June, 2020. SARS-CoV2 is an enveloped, positive-sense, single stranded RNA virus of genus Beta-coronavirus (β-CoVs) with the entire genome size of approximately 30kb. This viral genome is composed of about 14 open reading frames (ORF) which encodes both structural and non-structural proteins having a role in their transmission, survival and pathogenesis (1). The main structural proteins translated from sub-genomic mRNAs include spike glycoprotein (S), envelope protein (E), membrane glycoprotein (M) and nucleocapsid protein (N) along with 16 non-structural proteins (nsp 1-16). The genomic RNA element of SARS-CoV2 also includes 5’-untranslated region (5′-UTR) of 265bp length with a methylated cap and a 3’ polyadenylated UTR of length 229 bp, according to the reference sequence from Wuhan.

Low fidelity of the RNA polymerase makes the RNA viruses prone to high frequency mutation and the mutation determines the virus evolution (2). Systemic mutation analysis of the viral genome revealed that the virus had mutated several times in spatio-temporal variation and has evolved into numerous strains (3). This diversity of RNA strains might be correlated to severity and mortality seen in COVID-19 (Corona Virus disease-2019). A recent phylogenetic network analysis on 160 complete SARS-CoV2 genome reported that the virus appeared to evolve in three distinct clusters A, B and C, with A being the ancestral type. Type A and C seemed to be more prevalent in Americas and Europe, whereas type B was dominant in Eastern Asia (4).

During viral replication, subgenomic mRNAs are synthesized by a process of discontinuous transcription with a common 5′-untranslated leader sequence [5’-UTR] and a 3’-noncoding cotermini [3’UTR], identical to the viral genome (5). Hence, highly structured 3’ UTR of positive-strand RNA viruses is indispensable control element in replication, transcription, and translation of RNA viruses along with the 5’ UTR (6). Extent of structural and functional conservation in the 5’-terminal of genomic RNA of different species of genus coronavirus has been found to a distinctive magnitude (7). These terminal untranslated regions are thus substantial site for RNA-RNA interaction and binding of viral and host cellular proteins for RNA replication and translation (8).

Molecular evolution in the untranslated region i.e. the variation in the UTR region leads the virus to evolve to a great extent. There are many studies which have considered the mutation in the coding region with respect to geographical location. Here, we have enumerated the variants in the 5’- and 3’-UTR regions of SARS-CoV2 genome. We have studied a total of 8595 viral sequence samples worldwide for this purpose and found that there are some rare variants of UTR that are distributed over a global spectrum, while some variants are specifically present in a population at a comparatively higher frequency. This drove us to make a systemic catalogue of the UTR variants across six continents of the world that could have a role in emergence of different strains of SARS-CoV2. We have also looked at the possible regulation of viral genomic RNA through binding of host RNA binding proteins (RBPs) and miRNAs in specific sequences of the viral UTRs. There are experimentally validated evidences of human RBPs binding to the regulated signals within the untranslated region of SARS-CoV RNA in order to control the viral RNA synthesis and turnover. Polypyrimidine tract-binding protein (PTB) is found to bind to the leader sequence and *HNRNPA1, HNRNPQ* is bound to the 3’UTR of beta-coronavirus *MHV*.

Similarly, a host protein, *MADP1* (zinc finger CCHC-type and RNA binding motif 1) interacts with the 5′-UTR of SARS-CoV, influencing the RNA synthesis machinery (9). Likewise, there are reports of miRNA binding to the viral genome, both in the coding and non-coding regions (10). As the miRNA mediated decay of mRNA is observed mainly through the 3’UTR of the gene in mammalian system, we are also interested to explore the host miRNA-mediated regulation of the viral genome through the 3’-UTR of SARS-CoV2 via mRNA decay and translational repression. This interplay of host RBP and miRNA in the untranslated region of virus may delineate the role of UTRs in SARS-CoV2 virulence and survival and how variation in the UTR can have an impact in the overall regulation.

## Materials and Methods

### Viral genome sequences retrieval/resource

We have downloaded complete and high-coverage SARS-CoV2 genome sequence data in FASTA format from Global Initiative on Sharing All Influenza Data (GISAID), which is a public repository of all influenza virus sequence. Our data involved only the human-host specific viral sequence covering six continents of the world: Asia, Europe, Oceania, North America, South America and Africa. From Asia, we have taken the viral sequence data from China (n=197), India (n=205), Japan (n=123), Thailand (n=121) and Taiwan (n=102). From Europe, the data is taken from Italy (n=109), Spain (n=671), England (n=2345), Russia (n=130) and Germany (n=202). Viral sequence data is taken only from Australia (n=522), on behalf of Oceania. Representative countries from North America include USA (n=3286) and Canada (n=147). There are no further divisions for South America (n=281) and Africa (n=150). Initially the plan was to retrieve viral sequences collected in a span of one month, just after an interval of two month from the date of first report in the corresponding place, in order to catch the diversity in the viral sequence in that region. But, unfortunately, due to less number of sequence deposition in some countries, we had to deviate from the plan and retrieve all the available sequences till May28, 2020 for them (**Table 1**). We have downloaded the sequences on 28th May, 2020.

**Table 1:** Country/ continent wise list of number of SARS-CoV2 viral sequences used for this study

|  | **<u>COUNTRY</u>** | **<u>FIRST REPORT</u>** | **<u>DATA COLLECTION SPAN</u>** | **<u>NO. OF VIRUS SEQ. AVAILABLE & USED IN THIS STUDY</u>** |
| --- | --- | --- | --- | --- |
| **ASIA** | CHINA | 1-Dec-19 | Feb 02-March 01, 2020 | 197 |
|  | INDIA | 30-Jan-20 | April 01- April 30, 2020 | 205 |
|  | JAPAN | 3-Jan-20 | Data submitted till May 28, 2020 | 123 |
|  | THAILAND | 13-Jan-20 | Data submitted till May 28, 2020 | 121 |
|  | TAIWAN | 21-Jan-20 | Data submitted till May 28, 2020 | 102 |
| **EUROPE** | ITALY | 30-Jan-20 | Data submitted till May 28, 2020 | 109 |
|  | SPAIN | 31-Jan-20 | Data submitted till May 28, 2020 | 671 |
|  | ENGLAND | 31-Jan-20 | April 01- April 30, 2020 | 2345 |
|  | RUSSIA | 31-Jan-20 | April 01- April 30, 2020 | 130 |
|  | GERMANY | 27-Jan-20 | Data submitted till May 28, 2020 | 202 |
| **OCEANIA/**<br>**AUSTRALIA** |  | 25-Jan-20 | March 26- April 25, 2020 | 522 |
| **NORTH AMERICA** | USA/<br>PUERTO RICO | 20-Jan-20 | March 21- April 20, 2020 | 3286 |
|  | CANADA | 25-Jan-20 | Data submitted till May 28, 2020 | 147 |
| **SOUTH AMERICA** |  | 26-Feb-20 | February 26- Submitted till May 28, 2020 | 285 |
| **AFRICA** |  | 14-Feb-20 | February 14- Submitted till May 28, 2020 | 150 |
|  |  |  | **TOTAL** | **8595** |

### Alignment to reference sequence

The reference sequence from Wuhan was taken as our reference from NCBI (NC_045512.2). All the country specific sequences were aligned with the reference sequence separately using Multiple Sequence Alignment Tool Clustal Omega (EMBL-EBI). As we were interested only in the 5’- and 3’-untranslated region (UTR), and these positions are located at the two extreme ends of the sequences, there is a chance of lower coverage and higher error-rate. To resolve this ambiguity, we have put a high filtering cut-off of at least 70% coverage in a particular location of that area.

### Prediction of human RBP binding to viral UTR

For prediction of human RBPs bound to the UTR of a viral genome, we have used the web-tool ‘RBPmap’ that enabled prediction of RBP binding on genome sequences from a huge list of experimentally validated motifs of RBPmap database. Weighted-Rank algorithm was used for mapping the motifs (11). We have put the reference and mutated 5’- and 3’-UTR sequences separately as query sequence with the in-built human RBP motifs of the database. From the output, we have selected only the predicted RBPs with high z-score and p-value.

### Prediction of human miRNA interaction with viral UTR

Putative human miRNA binding sites on viral UTR was assessed using the ‘RNA22 v2 microRNA target detection’ tool (12). As a target sequence input, we provided the reference and mutated 3’-UTR sequences of virus genome. For input miRNA sequences, we have obtained a list of all annotated human miRNAs with their sequences in FASTA format from miRBase database (13). For all other criteria to be used for the prediction was given as per the default settings and additionally homology with seed sequence and its complementary target sequence were manually examined in the output file of interaction.

### Validation of expression of concerned RBP and miRNAs

Expression of specific genes was obtained from ‘The Protein Atlas’ respective to normal lung tissue. RNA-seq data from lung tissue generated by the Genotype-Tissue Expression (GTEx) project was reported as mean pTPM (protein-coding transcripts per million), corresponding to mean values of the different individual samples from each tissue. HPA RNA-seq data from lung tissue was reported as mean pTPM (protein-coding transcripts per million), corresponding to mean values of the different individual samples from each tissue. Tissue data for RNA expression also obtained through Cap Analysis of Gene Expression (CAGE) generated by the FANTOM5 project, reported as Scaled Tags Per Million. GEPIA (14) web-tool was used to obtain comparative expression of specific genes in lung adenocarcinoma (LUAD) and normal lungs.

## Results

### Worldwide cataloguing of prevalent UTR variants of SARS-CoV2

Sequence alignment of over 8500 virus samples collected from over 15 geographical locations with the SARS-CoV2 reference genome (NC_045512.2) has yielded a total of 74 variants in the 5’ UTR and 83 variants in the 3’UTR. Among them, 28 unique variants have been summarized in the **Supplementary Table 1** based on a minimum variant percentage of 0.5. But for the variants that are present in at least four populations, no such variant percentage cut off was taken. The most common high frequency variant is the 241C>T, which lies in the leader sequence of the SARS-CoV2 viral genome. China, being the ancestral domicile of the virus has a very low percentage of this variant (**Figure 1**). Europe has the maximum frequency of this variant (**Figure 2**). Other than China, other countries in Asia also have a lower percentage of this variant (**Figure 1**). Thus, we can a see a clear effect of regional variation on the frequency of this variant. Another study reported that this variant has co-evolved with three coding region mutations (3037C > T, 14408C > T, and 23403A > G) of nsp3, RNA primase and spike glycoprotein. They have also related this mutation with the increased transmissibility of the virus (15). Another common variation in the 5’UTR is 187A>G, that is present in seven of our populations being most prevalent in Italy and Canada. 29742G>A/T and 29774C>T are the most familiar variants of 3’UTR among all the continents. 29742 variant has two alternative alleles with G>A being more prevalent in India, USA and Africa, whereas G>T variant is more dominant in Thailand, Taiwan, Germany, Australia and Canada. China and England has both the variants of almost similar percentage. Hence, this variant seems not be associated with any specific region. Another important variant of 3’UTR is 29734G>C, which is mostly seen in Europe and Australia, but Italy has a quite high percentage of this variant than others (**Figure 2**). Most of the countries in this study seem to have a moderate to significant variation in the 3’UTR of the viral genome.

**Figure-1:**
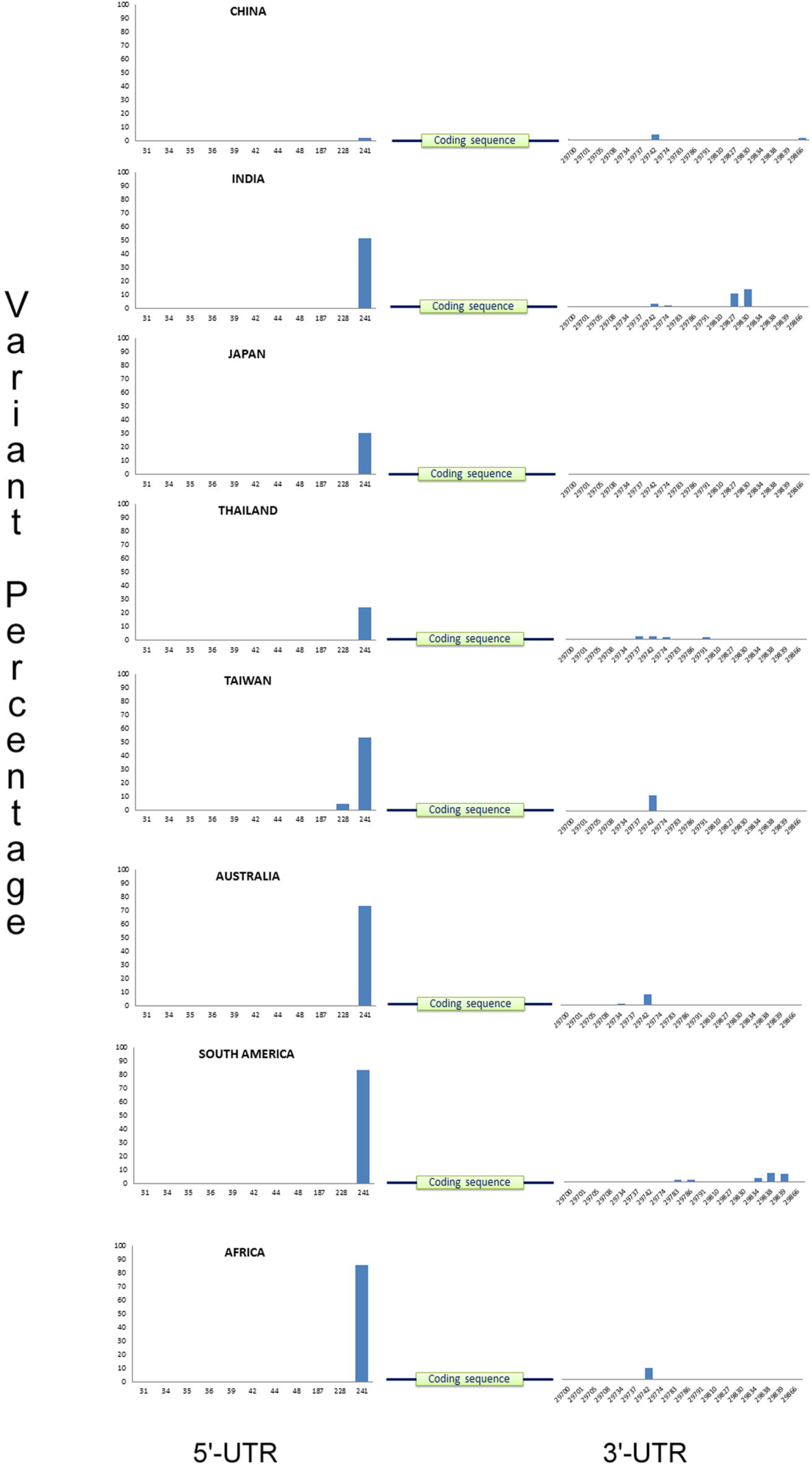
Global distribution of SARS-CoV2 Untranslated region variants from selected regions of Asia, Australia, South America and Africa. Both the 5’ and 3’-UTR variant positions are shown in ‘X’ axis and variant percentage has been shown in ‘Y’ axis.

**Figure-2:**
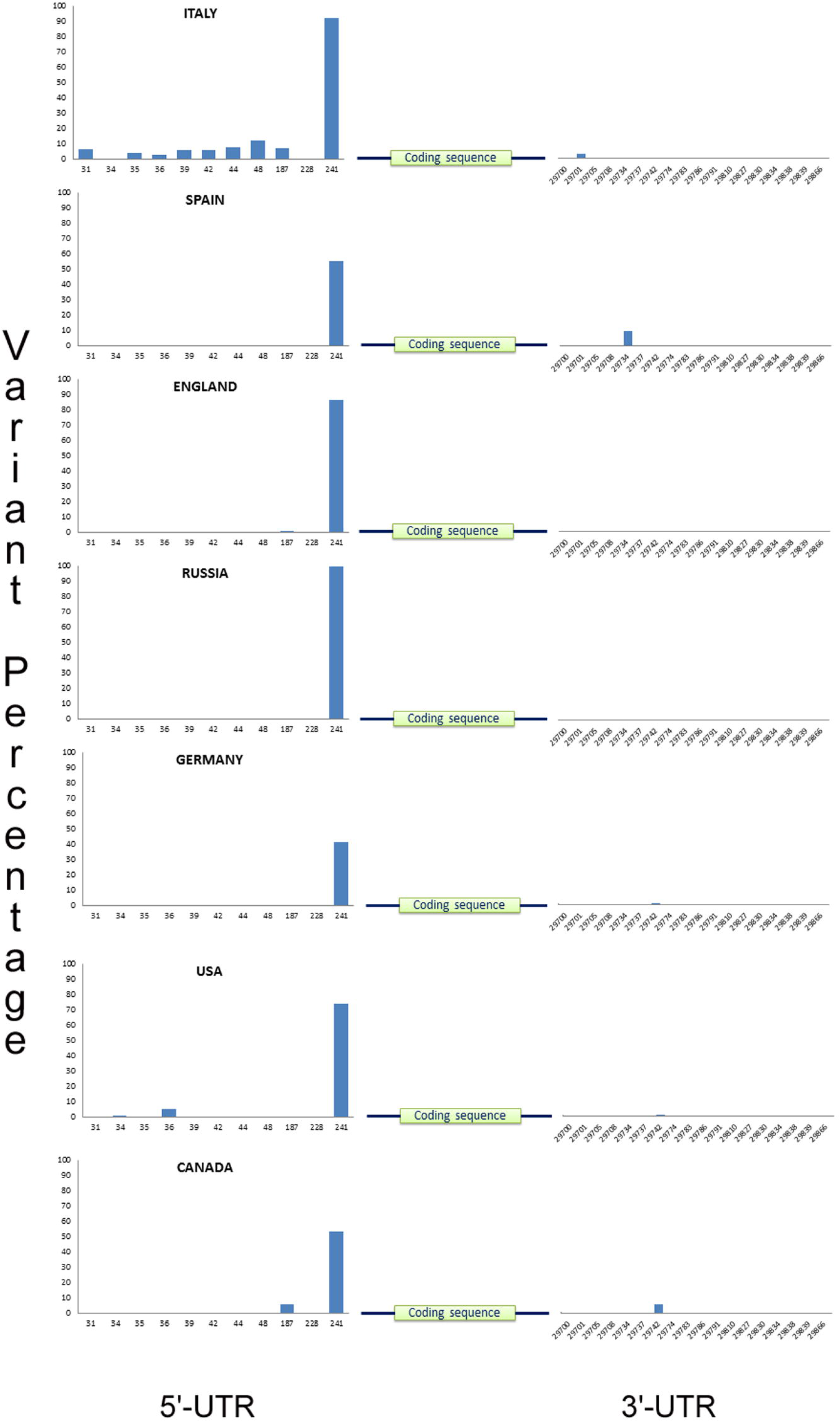
Global distribution of SARS-CoV2 Untranslated region variants from selected regions of Europe and North America. Both the 5’ and 3’-UTR variant positions are shown in ‘X’ axis and variant percentage has been shown in ‘Y’ axis.

### Country-specific UTR variant patterns

All the 15 populations and sub-populations used in this study, has a definite pattern of UTR variants of their own (**Figure 1; Figure 2**). Even countries in a single continent differ in their UTR variation. Some variations are merely specific to a particular location with a quite considerable percentage. The overall sequence of 5’UTR of SARS-CoV2 in Italy is exceptionally variable, whereas the 3’UTR is substantially stable with a single variant in the position 29701 (G>T). But England, also being a part of Europe, has a notable number of variations in the 3’UTR having a relatively stable 5’-end. From our analysis, it is evident that Russia had no such frequent variation in the UTR, except the 241C>T variant which was present in 100% of the population. India had two striking variations in the 3’UTR, 29827A>T, 29830G>T, which was found nowhere in this global mutational landscape (**Supplementary Table 1)**. All the sequences under this study with these two variants are from Indian state of Maharashtra, which has the highest number of SARS-CoV2 infected people within India. Most interestingly, both of this variants occur simultaneously in majority of the cases. Similarly, South America also had five unique mutations in the positions 29783G>T, 29786G>C, 29834T>C, 29838C>T and 29839A>G of 3’UTR.

### Identification of host RBPs bound to viral 5’-UTR

RNA binding proteins belong to a class of proteins which bind to target RNA molecules through characteristic binding motifs and perform a variety of functions. In fact, irrespective of the type of RNAs, whether is coding or non-coding, specific RBPs get associated with RNAs right after their birth and subsequently control the processing, stability, transport, translation, regulatory function and turnover. As mentioned earlier, exogeneous viral RNA will also encounter host RNA decay machinery when released into the cell and there are ample evidences that they attempt to escape the process in order to maintain their successful propagation within the host. One way to take care of the situation is to recruit host RNA stabilization factors into the regulatory region, i.e. 5’ and 3’-UTRs (16). **Table-2** lists the predicted host RBPs interacting with the 5’-UTR of virus SARS-CoV2 as obtained using ‘RBPmap’.

**Table 2:**
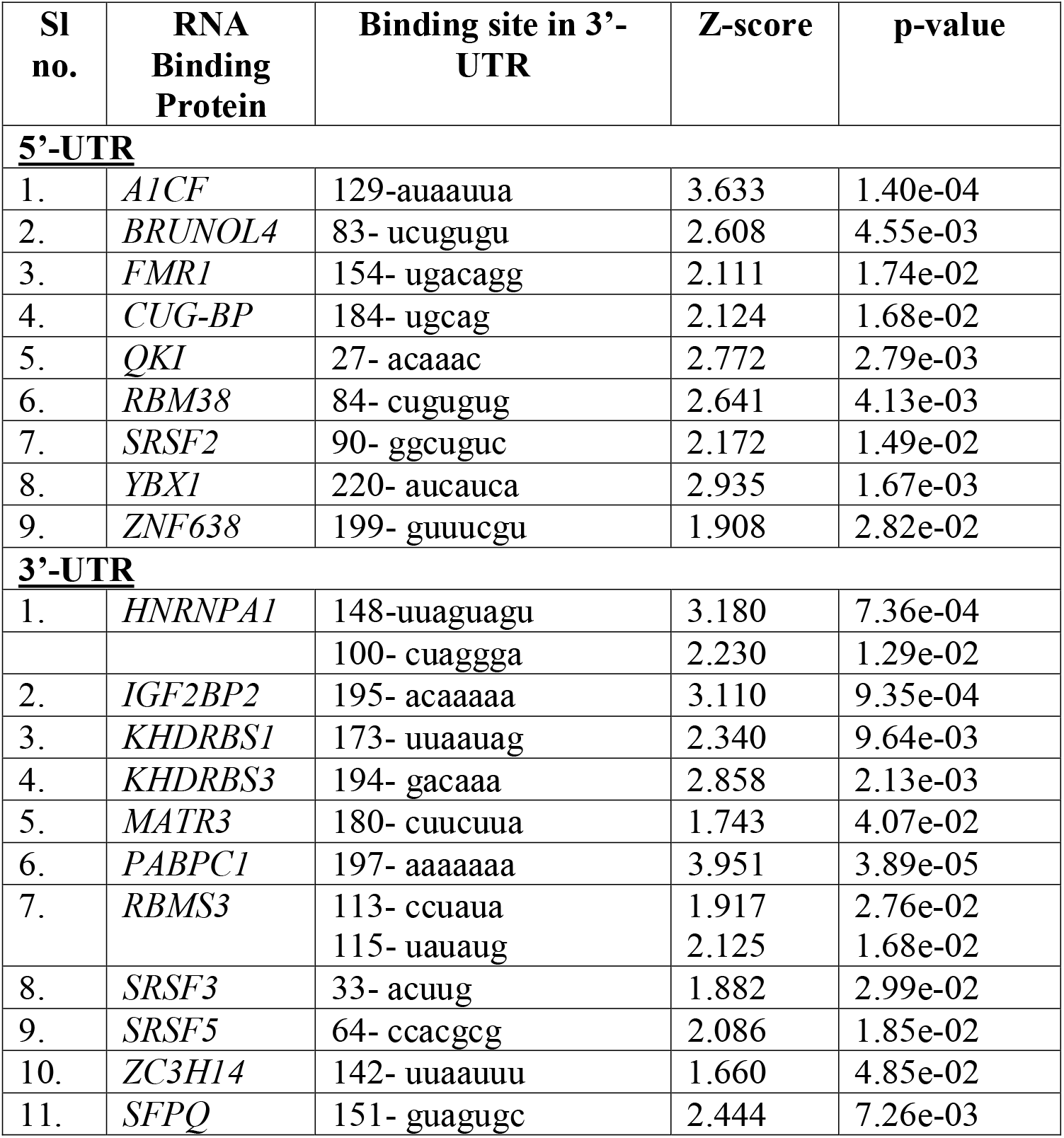
List of host RNA Binding Proteins as obtained from RBPmap, predicted to bind to 5’-UTR and 3’-UTR of SARS-CoV2 RNA

In the next step, the logical exploration was to find out whether the most prevalent variations identified in the 5’-UTR of the viral RNA genome could alter binding of any of these RBPs. The ‘A’ to ‘G’ change at position 187 coincided with the binding site of *CUG-BP* (also known as *CELF1*) (**Figure 3A**), implicating that the variation might have some effect on CUG-BP binding. In order to test that, we used the mutated sequence as ‘input’ and found that the single nucleotide change didn’t have any effect on binding of *CUG-BP* to its target sequence. In other words, no matter whether the virus has ‘A’ or ‘G’ at position 187 of its 5’-UTR, *CUG-BP* binds and probably performs its function.

**Figure-3:**
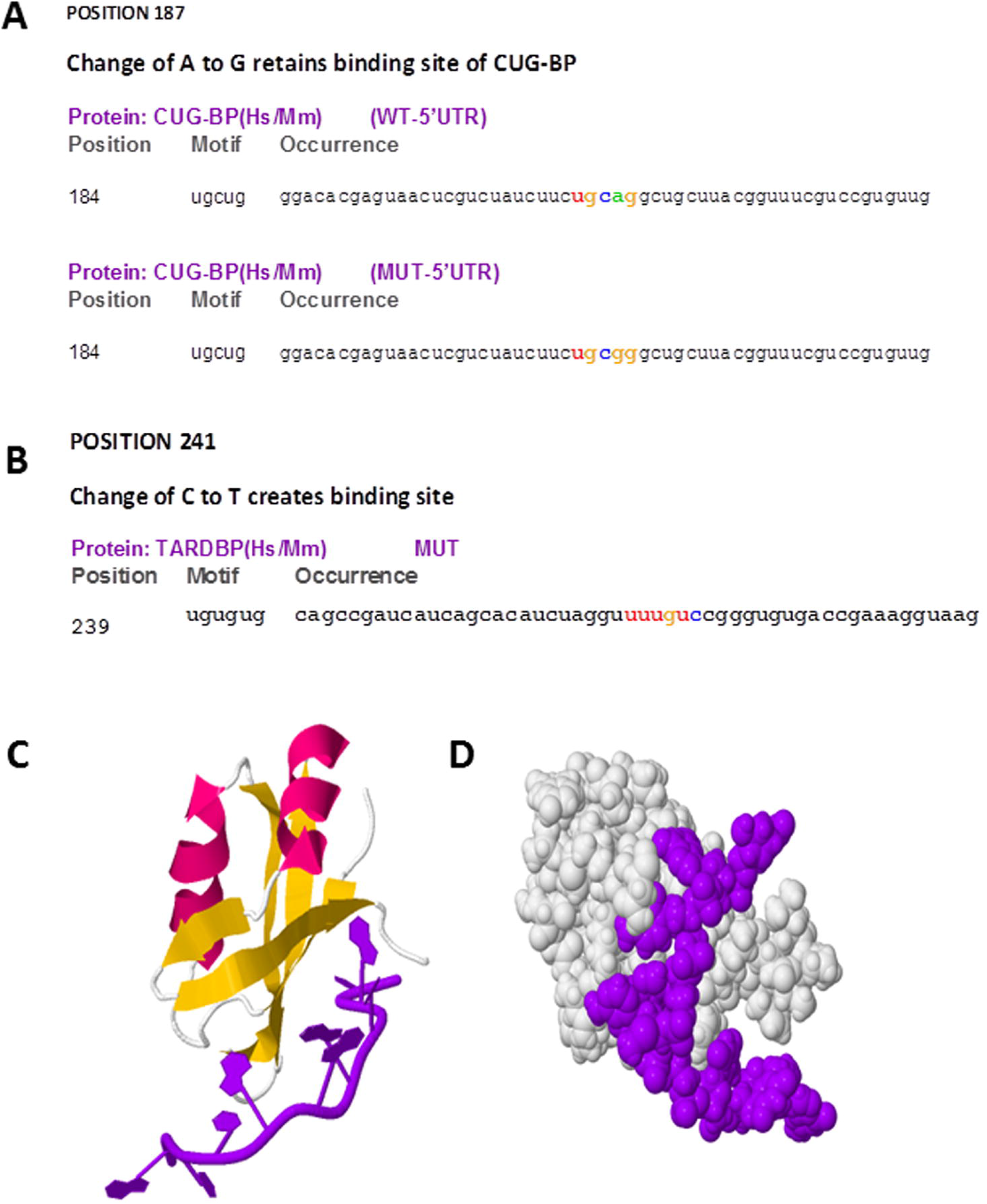
Effect of 5’-UTR variations on binding of RBPs at 5’-UTR. (A) Change of ‘A’ to ‘G’ at 5’-UTR position 187 retains binding site of CUG-BP (highlighted in coloured letters); while change of ‘C’ to ‘T’ at position 241 creates a binding site for TARDBP (highlighted in coloured letters) (B). Crystal structure of human TARDBP RRM1 domain in Complex with a single-stranded DNA (showed in ‘purple’) (PDB ID: 4IUF) in ribbon (C) and space filling (D) model, as retrieved from PDB.

Then, we focused on the most important change in the 5’-UTR, the position at 241. Looking at the distribution of this variation and the reports related to its association with virulence (15), we had a belief that it might be giving some selective advantage to the virus. However, we didn’t find any RBP binding site overlapping with this region. We changed the nucleotide at position 241 from ‘C’ to ‘T’ and used the mutated sequence in RBPmap. To our surprise, we found a *TARDBP* binding site created upon this change (**Figure 3B**). TARDBP (also called *TDP-43*) is a well characterized RBP which binds to specific ‘UG’ (and ‘TG’) rich sequences of RNA (and DNA) and reported to facilitate translation when bound to 5’-UTR (17). From protein data bank (PDB) we retrieved the co-crystal structure of TARDBP with single strand DNA (**Figure 3C & D**) which indicated strong binding of the protein to that particular nucleic acid stretch. Hence, our finding showed strong binding of *TARDBP* to viral 5’-UTR having ‘T’ at position 241. This could be implicated in facilitating translation of viral proteins resulting in its effective propagation within human host.

### Identification of host RBPs bound to 3’-UTR of SARS-CoV2

Following the same principle and considering the fact that 3’-UTR is the most important site for regulation of RNA stability and turnover, we also wanted to find out what are the RBPs interacting with this region. The 3’-UTR reference sequence of SARS-CoV2 was fed into RBPmap and we obtained a list of 3’-UTR binding RBPs selected on the basis of p-value and Z-score (**Table 2**). As with 5’-UTR, here also we focused on the major variations in 3’-UTR to find out if specific nucleotide change could interfere with the binding of specific RBPs. We found that 3’-UTR variations at positions 29742 and 29774 could affect interaction of *SRSF5* and *HNRNPA1* respectively. However, the regulation at 3’-UTR is not that simple. Another important factor which needs be taken into account is whether there are host miRNA binding sites present within the viral 3’-UTR region and how the combinatorial interaction of host RBPs and miRNAs could dictate the viral RNA stability.

### Finding of host miRNAs which could target viral 3’-UTR

In order to identify whether there are any host miRNA binding site present in viral 3’-UTR and if so, what they are, we used the viral 3’-UTR sequence as ‘input’ in the miRNA prediction software ‘RNA22 v2’. The algorithm predicts several of such miRNAs and we selected them based on folding energy for heteroduplex formation and p-value (**Table 3**). As the most important factor for the functional miRNA-target RNA interaction is to have the perfect homology between the seed sequence of miRNA (2^nd^ to 8^th^ nucleotide from 5’ end) and the corresponding target RNA sequence, we also explored the fact in the identified pairs. Furthermore, whether the changes caused by the common variations interfere with the seed sequence binding was also investigated. The interesting part in this scenario is that the reference sequence at a particular position causes a mismatch in the seed resulting in inefficient binding of a miRNA, but the variant base at that position creates a perfect binding site. However, considering the complexity of the region, all these probabilities could not be assessed only in terms of miRNAs and hence we set out to probe into the interaction between the host RBP-miRNAs and viral sequence variants in 3’-UTR region in a comprehensive manner.

**Table 3:** List of host microRNAs predicted by ‘RNA22 v2’ to have binding sites in the 3’-UTR of the viral genome

| Sl no. | miRNA | SARS-CoV2 3'-UTR Sequence corresponding to Seed region | folding energy (in -Kcal/mol) | p value |
| --- | --- | --- | --- | --- |
| 1. | hsa_miR_25_5p | 105-GGAGAGC | -15.40 | 3.3E-1 |
| 2. | hsa_miR_105_5p | 114-CTATATG | -13.20 | 3.3E-1 |
| 3. | hsa_miR_210_3p | 65-CACGCGGA | -17.80 | 2.24E-1 |
| 4. | hsa_miR_9_5p | 99-GCTAGGG | -14.82 | 3.3E-1 |
| 5. | hsa_miR_34b_5p | 110-GCTGCCT | -19.40 | 3.3E-1 |
| 6. | hsa_miR_196b_5p | 109-GCTGCCT | -16.60 | 3.3E-1 |
| 7. | hsa_miR_1293 | 63-GCCACGC | -19.90 | 2.24E-1 |
| 8. | hsa_miR_2116_5p | 121-GAAGAGC | -14.90 | 3.3E-1 |
| 9. | hsa_miR_323b_5p | 105-GAGAGCT | -19.70 | 3.3E-1 |
| 10. | hsa_miR_4497 | 66-ACGCGGA | -16.10 | 2.24E-1 |
| 11. | hsa_miR_4659b_3p | 120-GGAAGAG | -18.80 | 3.3E-1 |
| 12. | hsa_miR_4757_5p | 59-CGAGGCC | -20.90 | 2.24E-1 |

### Interaction between RBPs and miRNAs

We investigated the interaction in a stepwise manner. The first situation explored was the case where host miRNA miR-34b-5p targets viral RNA and no mutation has been identified in the ‘seed’-corresponding region. This means that all the viral sub-types will be vulnerable to degradation by this miRNA, provided it is expressed in host lungs tissue. Careful scanning of the binding site and also looking at the RBP binding information we identified a *RBMS3* target sequence actually overlapping with the miR-34b-5p seed corresponding region (**Figure 4A**). This clearly indicated a prevalent competition between *RBMS3* and miR-34b-5p-RISC complex for their respective binding site, with one of them simply presenting a steric hindrance to the other one by blocking the access. This is quite common scenario in regulation of mammalian mRNA stability where the final verdict whether the mRNA is degraded or stabilized largely depends on the spatio-temporal expression or the availability of both the molecules (RBP and miRNA).

**Figure-4:**
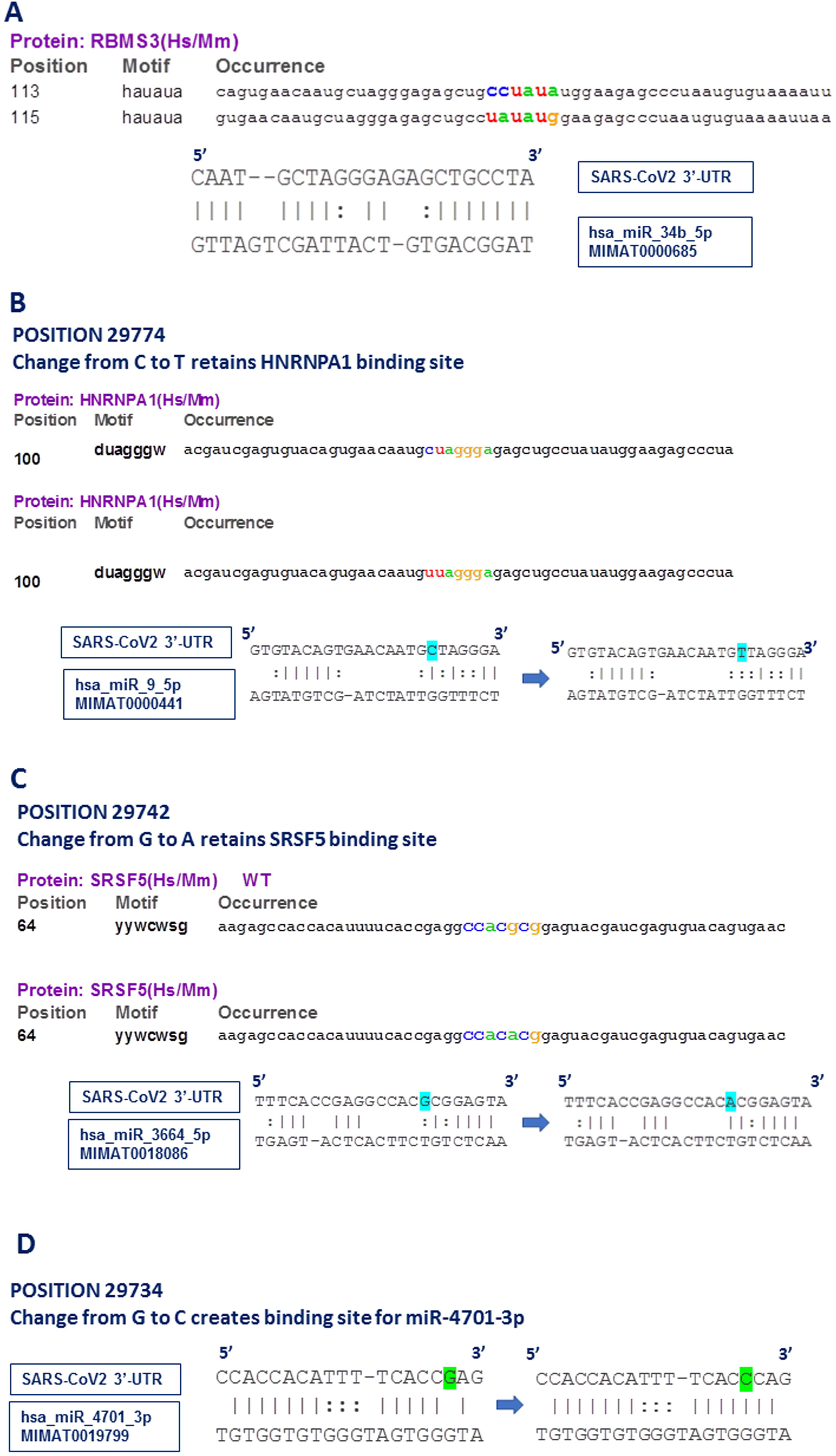
Interaction between RBPs and miRNAs at 3’-UTR. (A) RBMS3 and miR-34b-5p binding site overlap with each other. RBMS3 binding site highlighted in coloured letters. (B) Change from ‘C’ to ‘T’ at position 29774 shows intact binding site for HNRNPA1 (highlighted in coloured letters) and disrupted interaction of miR-9-5p with its target (sequence change highlighted in blue). (C) Change from ‘G’ to ‘A’ at position 29742 shows intact binding site for SRSF5 (highlighted in coloured letters) and creation of binding site of miR-3664-5p with its target (sequence change highlighted in blue). (D) Change from ‘G’ to ‘C’ at position 29734 shows newly created interaction of miR-4701-3p with its target (sequence change highlighted in green).

Second scenario was the variation in position 29774. In this case, we identified that change from ‘C’ to ‘A’ disrupted the binding of miR-9-5p to its corresponding target sequence at the ‘seed’ region (**Figure 4B**). This is clearly an advantage to the virus as the variation would help it to get rid of the miRNA mediated degradation. As this overlapping region also had a binding site for RBP *HNRNPA1*, we wanted to find out what happens to *HNRNPA1* binding when there is a change from ‘C’ to ‘A’. Interestingly enough, we discovered that HNRNPA1 binding is also not affected, implicating an ensured protection of viral RNA by RBP *HNRNPA1*, no matter whether miR-9-5p is functional or not. Again, this situation is also quite common in mammalian mRNA regulation where we see some amount of degeneracy among the RBP target sequences, when change of a single nucleotide is often tolerated. The virus uses this phenomenon to its benefit where despite of UTR sequence variation protection from same RBP is ensured.

In case of the third incidence, we looked at position 29742, where the ‘G’ to ‘A’ change creates a binding site for miR-3664-5p (**Figure 4C**), which means that the virus having the variant nucleotide will be prone to degradation by this miRNA. We had already commented about the possible binding of *SRSF5* in this region and found that the variation under investigation had no impact on *SRSF5* binding. Again, it indicated that despite of creation of miR-3664-5p binding site, the viral RNA is safe until *SRSF5* is available to bind and compete with miR-3664-5p-RISC complex. However, as evident from our 3’-UTR analysis, a significant proportion of viral sequence had ‘G’ to ‘T’ change at position 29742. While *SRSF5* binding site was still retained, the miR-3664-5p site creation did not take place due to this change.

The fourth scenario was found to be very different from what have been mentioned so far. We identified that change of ‘G’ to ‘C’ at position 29734 creates a binding site for miR-4701-3p, but no RBP with a overlapping target sequence could be identified (**Figure 4D**). This means that there is probably no host factor to protect viral RNA from miR-4701-3p mediated degradation when the virus carries a ‘C’ at position 29734. However, this entirely depends on the expression status of miR-4701-3p in host tissue and there are evidences that suppression of miRNA expression falls under viral strategy to safeguard its genome.

### Validation of our finding by checking expression patterns of key RBPs and miRNAs in host tissue

The most important aspect of our study lies in the fact how we could validate our finding. The phenomenon of viral RNA utilizing host stabilizing factors and escaping RNA decay machinery is well-established. But, nothing is known so far for SARS-CoV2. As mentioned before, the actual incident what is taking place after viral infection largely depends on the expression status of the interacting RBPs and miRNAs in particular host tissue. Since, lungs is the primary site of infection for SARS-CoV2, we wanted to find out what is the nature of expression of these molecules in normal lungs tissue. In the first approach we tested their expression in ‘The Protein Atlas’ portal (**Figure 5 A, B**) and in the second approach we used the TCGA dataset for lung adenocarcinoma (LUAD) from GEPIA and compared the expression of the selected RBPs in adjacent normal lung tissue (**Figure 5 C, D**). To our excitement, we found a decent to huge expression of all the RBPs in normal lung tissues, clearly indicating their availability to protect viral RNA from host decay machinery. As the viral infection will follow a huge inflammatory condition in the lungs and by the time we know that in severe cases this prolonged inflammation leads to pulmonary fibrosis, we also wanted to check whether there are supporting evidences relating the expression of these RBPs and miRNAs to inflammation or more specific to pulmonary inflammation. As shown in **Table 4**, almost all the RBPs under investigation were either reported to have induced by inflammation or responsible for fibrosis in lungs or other organ. Expression of some of them was also reported to be modulated by viral infection. This finding supports our hypothesis and indicates that from the stages of early viral infection to severe advanced conditions SARS-CoV2 could utilize the host stabilization factors for its own benefit.

**Figure-5:**
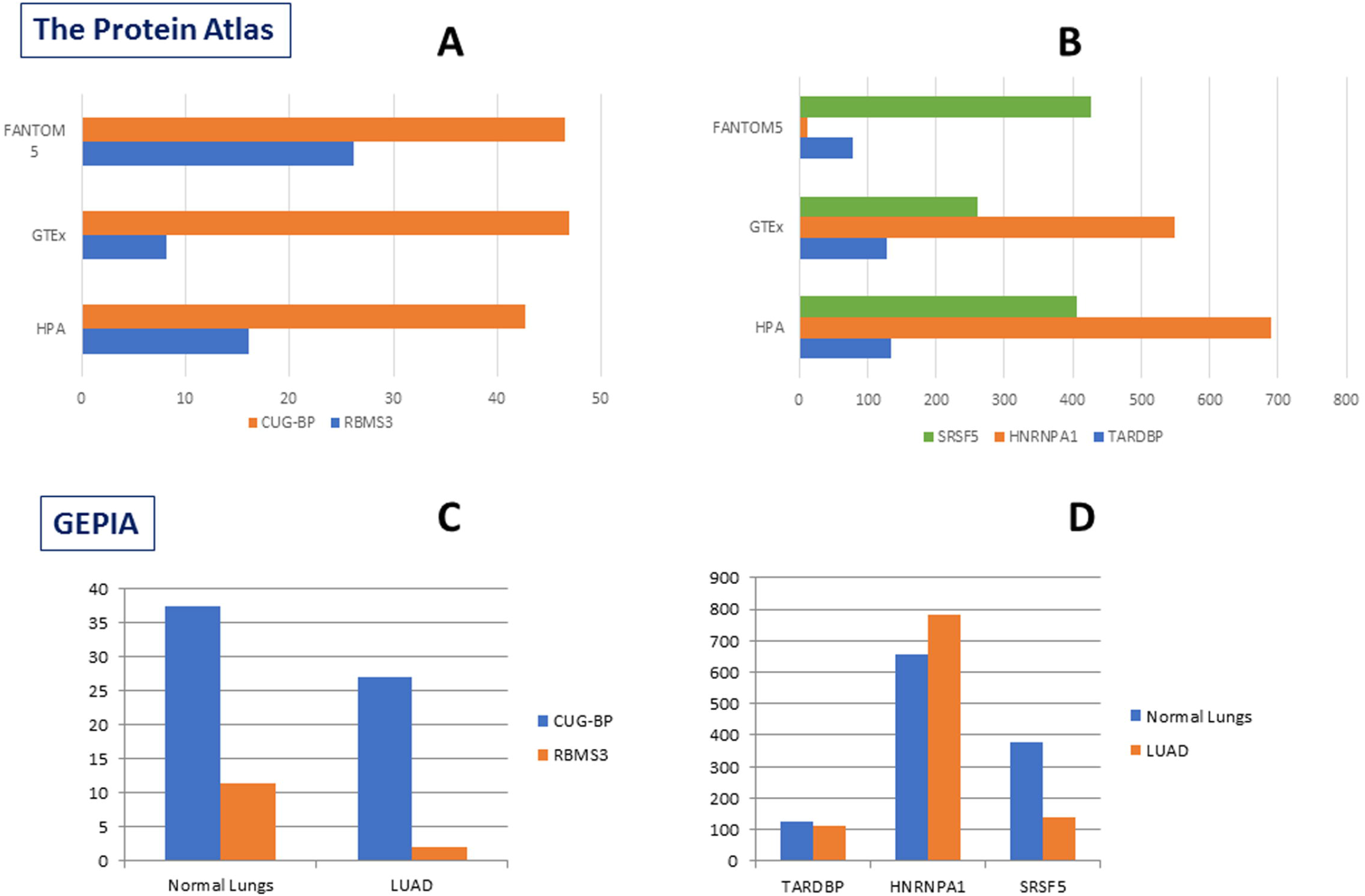
Expression of selected RBPs in lungs tissue. (A, B) Expression of five selected RBPs in Lungs tissue of normal individuals as obtained from ‘The Protein Atlas’. RNA-seq tissue data generated by GTEx, HPA and FANTOM5 projects showed expression pattern of these genes in normal lungs. (C, D) Expression of five selected RBPs in Lungs adenocarcinoma (LUAD) and adjacent normal lungs tissue as obtained from ‘GEPIA’.

**Table 4:**
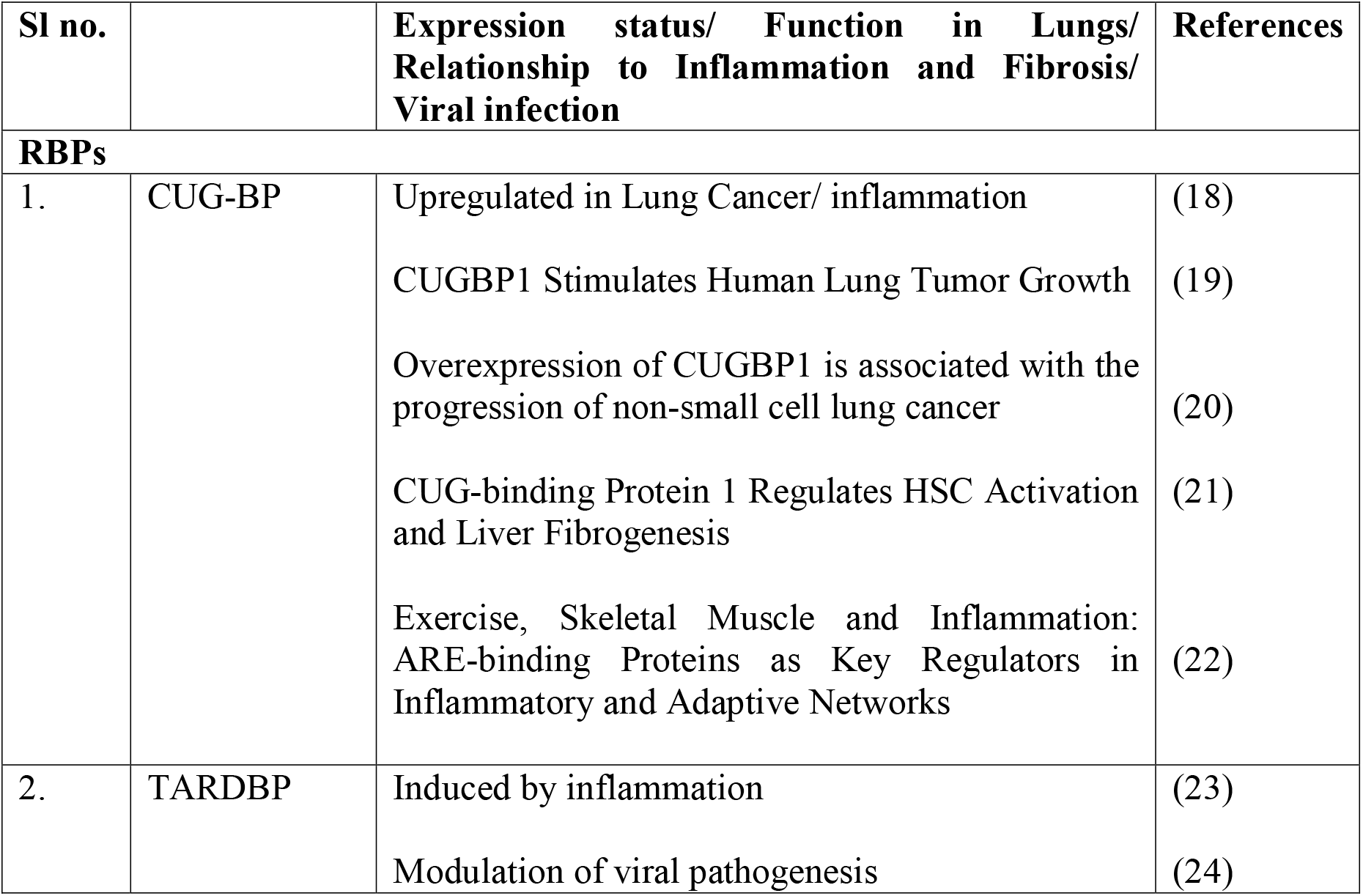
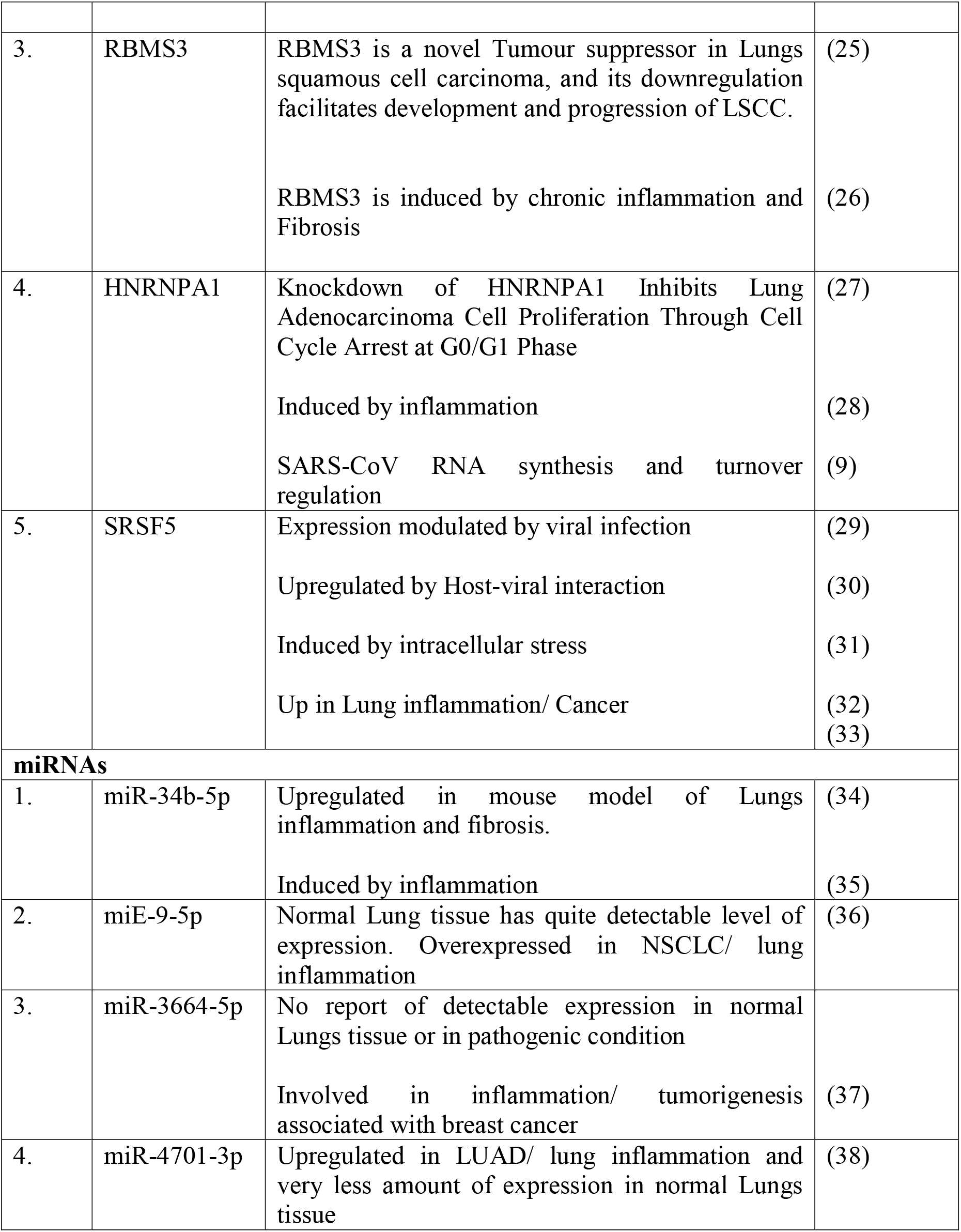
List of supporting experimental evidences validating our findings

Following the same tune, we looked at the miRNAs also in greater details and several interesting observations were noted. Apart from miR-3664-5p, other three miRNAs were detectable in lungs tissue and even were closely associated with pulmonary inflammation. In case of miR-3664-5p also, its involvement in breast cancer tumorigenesis and associated inflammation was quite supportive. Most of these supporting studies mentioned here for both RBPs and miRNAs are actually functional interrogations involving rigorous experimental procedures. Hence, we are quite confident that that our predictions got validated by experimental reports asking the similar type of questions in a similar set up.

## Discussion

It has been more than six months we are experiencing one of the biggest pandemics of the world. Investigation of the viral subtypes across the globe has identified several variants in the coding region of the viral RNA possibly resulting in key structural or functional changes in spike protein, nucleocapsid proteins or the RNA dependent RNA polymerase of the virus. As compared to the coding region variants, there have not been so many changes identified in the 5’ or 3’ untranslated regions of the viral genome. This made us interested about these regions and first we wanted to carry out a systematic exploration of the prevalent changes present in the region using more than 8500 viral sequences isolated from all the continents and explore their significance with respect to host RNA decay machinery as well. Despite of some specific changes concentrated in some specific regions of the world, overall pattern of the 5’ and 3’-UTR variants pinpoint on few predominant ones. The most important of it was at position 241, where we see a clear change of distribution of reference to variant nucleotide from China to South-East Asia to Europe and Americas. This change has also been implicated to higher mortality and/ or infectivity of the virus (15). For the first time, we report the likelihood of TARDBP binding to this region. *TARDBP* has been known to promote translation and RNA stability and it has also been shown to play important role in viral infection (39, 40). Therefore, promotion of translation of SARS-CoV2 viral proteins by *TARDBP* fits well with possible selection of ‘T’ base over time and its correlation to infectivity.

The other RNA binding proteins identified in the study to be interacting with viral 5’ and 3’-UTR also has reported evidences of functioning as RNA stabilizing factor or facilitator of translation. *CUG-BP* has been shown to interact with *eIF2A* (41) and promote translation in selected targets (42). *RBMS3* is a very well-characterized RNA stabilizing factor and known to increase half-life of target mRNAs as well as increase protein expression in many instances (26). Similarly, *HNRNPA1* is also a RBP known for its role in promoting mRNA stability after binding to 3’-UTR specific target sites and as mentioned before, this RBP has been experimentally identified to bind to SARS-CoV 3’-UTR (43, 44). SRSF5, although primarily involved in splicing, has been attributed to functions like maintaining mRNA stability and translation (45, 46).

The most significant aspect of this study was to identify miRNA binding site in the viral 3’-UTR and deciphering the cross-talk between RBPs and miRNAs along with the nucleotide variations at specific sites. Our results elaborated five distinct types of such interactions, as summarized in **Figure 6**. We have seen *CUG-BP* binding site is retained and TARDBP binding site is created upon change in viral nucleotide sequences at target positions of 5’-UTR, providing a continued stability and translational advantage to the virus (shown in arm:A). On the other hand, there exists multiple possibilities in case of 3’-UTR, where we first see direct competition over access to target sites between *RBMS3* and miR-34b-5p (arm:B). Lung inflammation causes induction of miR-34b-5p (34) which could be a step to attenuate viral infection. However, the sustained expression of RBMS3 in lungs tissue probably acts to prevent miR-34b-5p action after binding to the viral RNA at overlapping site.

**Figure-6:**
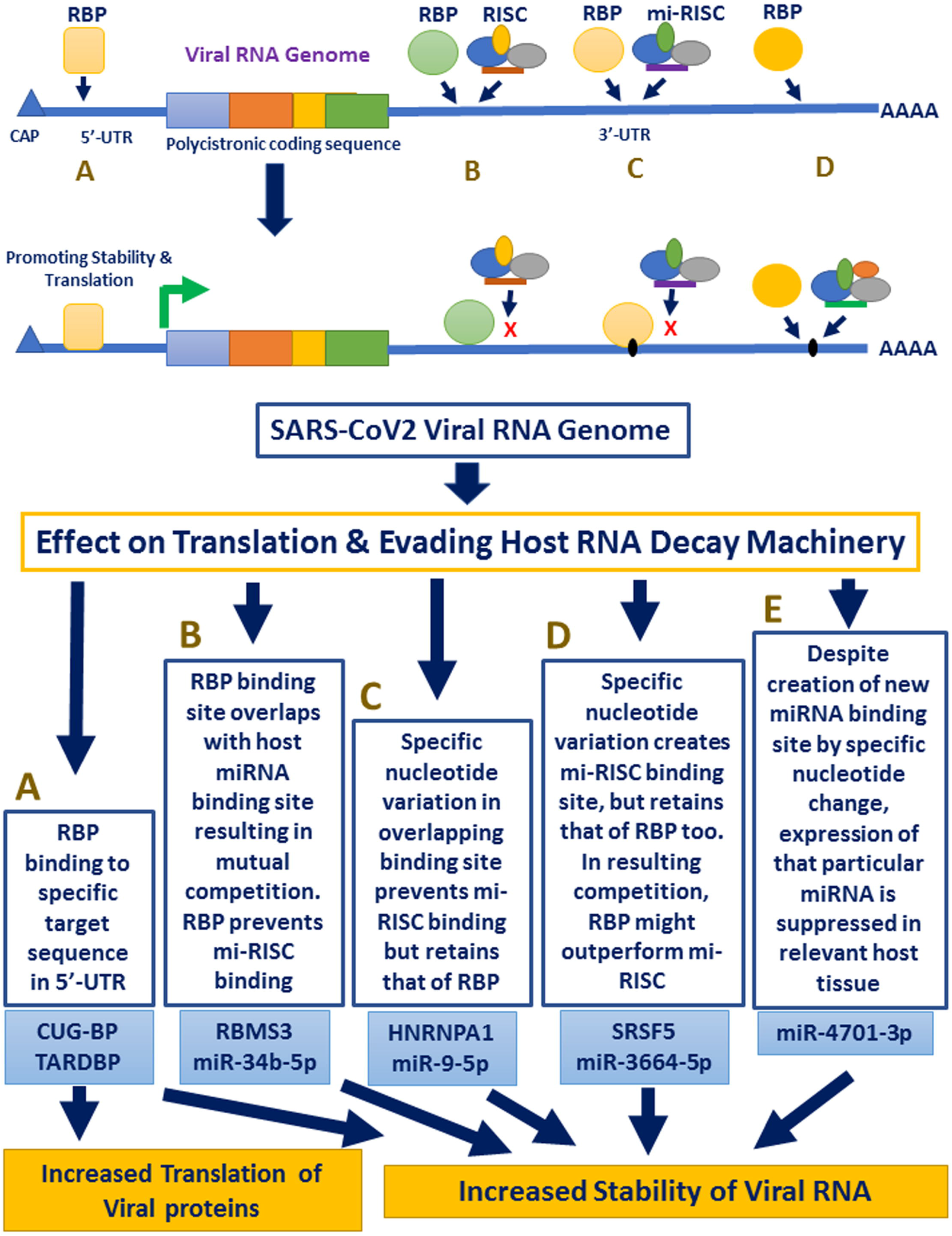
Schematic representation of host RBP-miRNA interactions along with sequence variation in UTR regions of SARS-CoV2.

The effect of variation in viral sequence could either disrupt or create miRNA binding site, keeping the RBP binding site intact. This result in non-functional miRNA site in one case (arm:C) and competition between RBP and miRNA in the other (arm: D). Expression of both *HNRNPA1* and *SRSF5* is high in lungs, further supporting their possible protective effect on viral genome. The incidence described in arm: E is different as it creates a miRNA binding site without an RBP protection. We have seen that expression of miR-4701-3p is less in lungs, provoking the thought that despite having its site created, there might not be enough miRNA to act on the viral RNA at this site.

Thereby, we, for the first time, characterized SARS-CoV2 5’ and 3’-UTR sequences for existing variations in these regions prevailing at major countries of infection spread over all the continents. We further identified interactions between host RBPs like *CUG-BP, TARDBP, RBMS3, HNRNPA1* and *SRSF5* with host miRNAs miR-34b-5p, miR-9-5p, miR-3664-5p and miR-4701-3p and showed how the interactions changed along with sequence variations at specific positions of untranslated regions of viral genome. Our findings elaborate complex relationship between host RNA stabilization/ decay factors and SARS-CoV2 RNA genome highlighting how the virus could manipulate host machinery. The knowledge could also be used to develop antiviral compounds following further experimental studies.

## Acknowledgements

We acknowledge the support and advice of Prof. Saumitra Das, Director, NIBMG and time to time suggestions from Dr. Bornali Bhattacharjee and Dr. Bhaswati Pandit, NIBMG. MM received fellowship from University Grants Commission, Government of India.

## List of abbreviations

SARS-CoV2: Severe Acute Respiratory Syndrome Coronavirus 2
miRNA: microRNA
nsp: Non-structural protein
HNRNPA1: Heterogeneous Nuclear Ribonucleoprotein A1
HNRNPQ: Synaptotagmin Binding Cytoplasmic RNA Interacting Protein
MHV: Mouse Hepatitis Virus
MADP1: Zinc Finger CCHC-Type And RNA Binding Motif Containing 1
EMBL-EBI: European Molecular Biology Laboratory-European Bioinformatics Institute
GEPIA: Gene Expression Profiling Interactive analysis
*CUG-BP*: CUG rich region binding protein
*CELF1*: CUGBP Elav-Like Family Member 1
*TARDBP*: TAR DNA Binding Protein
*SRSF5*: Serine And Arginine Rich Splicing Factor 5
*RBMS3*: RNA Binding Motif Single Stranded Interacting Protein 3
TCGA: The Cancer Genome Atlas
*eIF2A*: Eukaryotic Translation Initiation Factor 2A

## Declarations

### Ethics approval and consent to participate

The study does not involve any patient sample, but uses only sequences shared in public domain. Hence, not applicable.

### Consent for publication

Not applicable

### Availability of data and materials

Not applicable

### Authors’ contributions

MM and SG carried out the sequence analysis and RBP-miRNA analysis. SG conceptualized, designed the study. Both MM and SG drafted the manuscript, prepared and approved the final version.

### Pre-print information

The manuscript has been submitted in pre-print server bioRxiv as: **doi:** https://doi.org/10.1101/2020.06.09.134585

---

**Supplementary Table 1**: List of variations in 5’ and 3’-UTR of SARS-CoV2 across six continents of the world.

